# Transcription-coupled DNA repair underlies variation in persister awakening and the emergence of resistance

**DOI:** 10.1101/2021.07.29.454265

**Authors:** Wilmaerts Dorien, Focant Charline, Matthay Paul, Michiels Jan

## Abstract

Persisters constitute a population of temporarily antibiotic-tolerant variants in an isogenic bacterial population and are considered an important cause of relapsing infections. It is currently unclear how cellular damage inflicted by antibiotic action is reversed upon persister state exit and how this relates to antibiotic resistance development. We demonstrate that persisters, upon fluoroquinolone treatment, accumulate oxidative damage which is repaired through nucleotide excision repair. Detection of the damage occurs via transcription-coupled repair using UvrD-mediated backtracking or Mfd-mediated displacement of the RNA polymerase. This competition results in heterogeneity in persister awakening lags. Most persisters repair the oxidative DNA damage, displaying a mutation rate equal to the untreated population. However, the promutagenic factor Mfd increases the mutation rate in a persister subpopulation. Our data provide in-depth insight in the molecular mechanisms underlying persister survival and pinpoints Mfd as an important molecular factor linking persistence to resistance development.

## Introduction

Every isogenic bacterial population harbors a subpopulation of non-growing antibiotic-tolerant cells that survive lethal drug exposure, named persisters. Upon antibiotic removal, a persister cell that resumes growth can form a new bacterial population. Persistence is an example of phenotypic bistability leading to the presence of antibiotic-sensitive and -tolerant cells in an isogenic population (1, 2). Evidence for the clinical importance of persister cells is accumulating, as they are held responsible for the recalcitrance of chronic infections (3–6) and they constitute a pool from which resistant mutants emerge (7–10). Indeed, antibiotic-resistant mutants emerged from antibiotic-tolerant cells in *Escherichia coli, Mycobacterium tuberculosis* and *Staphylococcus aureus* (8–10). Furthermore, a positive correlation was found between the number of persisters in a bacterial population, and the emergence of resistant mutants in lab and natural *E. coli* strains (7). In this respect, the healthcare system would benefit from the development of anti-persister therapies.

The number of persisters at any given time is determined by a combination of the rates of persister state entry and exit. Several decades of persistence research have resulted in the discovery of different molecular mechanisms triggering persister state entry (1). These mechanisms often include decreased cellular energy levels (11, 12) and/or a block in essential cellular processes such as translation (13, 14) and replication (15). In addition, mechanisms that avoid antibiotic-inflicted DNA damage or antibiotic accumulation in the cell were reported to induce the persister state (16, 17). In stark contrast, only few studies have attempted to investigate mechanisms coordinating persister state exit. This discrepancy mainly results from the technical difficulties in analyzing and quantifying persister awakening, such as the transient phenotype and the small number of persisters in a population. As a result, a systematic understanding of persister state exit is largely lacking.

Recent work has postulated the hypothesis that persister state exit depends on reverting the physiological changes correlated with the induction of the persister state (1). For example, it has been hypothesized that persisters formed by the TacT toxin, which acetylates tRNA, resume growth via Pth-mediated deacetylation (14). Additionally, persisters formed by the pore-forming HokB toxin were found to wake up based on pore disassembly via a combination of DsbC-mediated monomerization and DegQ-mediated peptide degradation, followed by membrane repolarization (18). Furthermore, protein aggregate formation and removal were shown to be correlated with persister state entry and exit (19). Based on these findings, we can infer that other persister formation pathways should also be reverted to initiate persister state exit.

It has long been assumed that persisters have a dormant phenotype which allows them to survive the antibiotic treatment because of target inactivity (20). However, recent work has pointed towards a role for antibiotic-induced damage repair in successful persister awakening. Indeed, both persisters from cultures in stationary or exponential phase were found to acquire DNA-damage after fluoroquinolone treatment (21, 22), contradicting this hypothesis. Later, it was demonstrated that the timing of DNA-damage repair is important for successful persister recovery after antibiotic treatment by DNA-damaging agents. Indeed, if DNA repair was stalled until the initiation of growth-associated processes, persister survival was strongly reduced (23). Furthermore, ploidy was found to enhance persister survival, demonstrating that persisters use homologous recombination (HR) to repair the double-stranded breaks (DSB) (24). Combined, DNA damage repair has proven to be an essential and general phenotype for persister survival following fluoroquinolone treatment, making it a promising target for an anti-persister therapy. However, research to date has not identified any molecular mechanisms that underlie the successful recovery of the persister cell. To address which of these repair proteins are important for successful persister recovery, we systematically tested deletion mutants of DNA-repair genes and examined their expression during recovery following antibiotic treatment.

In this work, five DNA-damage repair genes (*recB, recC, uvrA, uvrB* and *uvrD*) were found to be important for persister survival following fluoroquinolone treatment. RecB and RecC are both involved in double strand break repair via HR (25). The UvrA2B complex, together with UvrC and UvrD, is involved in nucleotide excision repair (NER) (26). NER repairs a broad variety of bulky, structurally unrelated, DNA damages, such as DNA damage following UV irradiation or hydrogen peroxide treatment (27, 28). Damage recognition is based on either UvrA2B that scans the entire genome and detects damages that cause a distortion of the double helix (26) or via transcription-coupled repair (TCR), which is based on the detection of DNA lesions that completely stall (29, 30) or pause (31) the RNA polymerase, after which repair is orchestrated by NER. TCR is biased towards actively expressed genes (30, 32). The bacterial cell employs two different TCR pathways. The first one relies on Mfd, which recruits UvrA to the site of the lesion for NER-mediated repair and pushes the RNA polymerase forward, terminating transcription (33, 34). The translocase Mfd is also an evolvability factor and was found to increase mutagenesis (35, 36). A second TCR pathway was found to depend on UvrD, which backtracks the stalled RNA polymerase, after which NER-mediated repair occurs following NusA-mediated recruitment (37). The RNA polymerase is not lost upon backtracking by UvrD, enabling transcription to resume rapidly upon DNA repair (37). Presence of (p)ppGpp makes the RNA polymerase backtracking-prone (38). After detection of the DNA lesion, UvrC cleaves 5’ and 3’ from the lesion, after which the helicase UvrD releases the damaged oligonucleotides (26, 39). In addition to its role in NER and TCR, UvrD acts as a helicase in methyl-directed mismatch repair following replication (40) and helps stalled replication forks to restart by removing the Tus protein (41, 42).

In this work, we demonstrate that ofloxacin treatment results in the accumulation of oxidative DNA damage. Transcription re-initiates during recovery in fresh medium, enabling TCR to detect and repair the oxidative DNA damage via NER. Clearing of the RNA polymerase from the site of the lesion occurs via UvrD or Mfd. Single-cell observations of recovering persister cells demonstrate that the UvrD-mediated TCR pathway decreases the awakening lag of the persister cell, while Mfd-mediated TCR increases the awakening lag. Furthermore, Mfd-mediated TCR is responsible for the emergence of a subpopulation of persister cells with a high mutation rate, providing mechanistic insight on the emergence of resistant mutants from a persister population. In addition, phenotypic characteristics of persister cells were compared with sensitive cells, revealing differences in initial elongation rate during the awakening lag and in *mfd* expression levels. Combined, our work is one of the first to explore molecular mechanisms underlying persister awakening dynamics in a wild-type background and provides us with an understanding of the accumulation of mutations in persister cells at the molecular level.

## Experimental work

### 1. Absence of RecB, RecC, UvrA, UvrB and UvrD impedes persister awakening after fluoroquinolone treatment

To define which DNA repair pathways underlie persister cell survival after fluoroquinolone treatment, we hypothesized that absence of a gene important for survival should result in a decreased persister fraction. Therefore, we have first built a comprehensive list of genes coding for proteins that are directly implicated in DNA damage repair (Table S1). Next, the persister fractions of the corresponding deletion mutants were assessed in stationary phase after treatment with the fluoroquinolone antibiotic ofloxacin. Figure 1A demonstrates that absence of *recB, recC, uvrA, uvrB* and *uvrD* results in a strong decrease in the persister fraction following ofloxacin treatment. Time-kill curves were performed to confirm the persister population (Fig. S1A) (43). The lower persister fraction of *recB* and *recC* mutants nicely demonstrates the validity of our hypothesis, as RecB and RecC were recently linked with persister survival following their role in homologous recombination to repair DSB (24). UvrA, UvrB and UvrD are components of the NER pathway, and the molecular mechanisms underlying their role in persistence are unknown (23, 26, 44). We confirmed the decreased survival of these mutants following treatment with the fluoroquinolone moxifloxacin (Fig. S1B).

**Figure 1.**
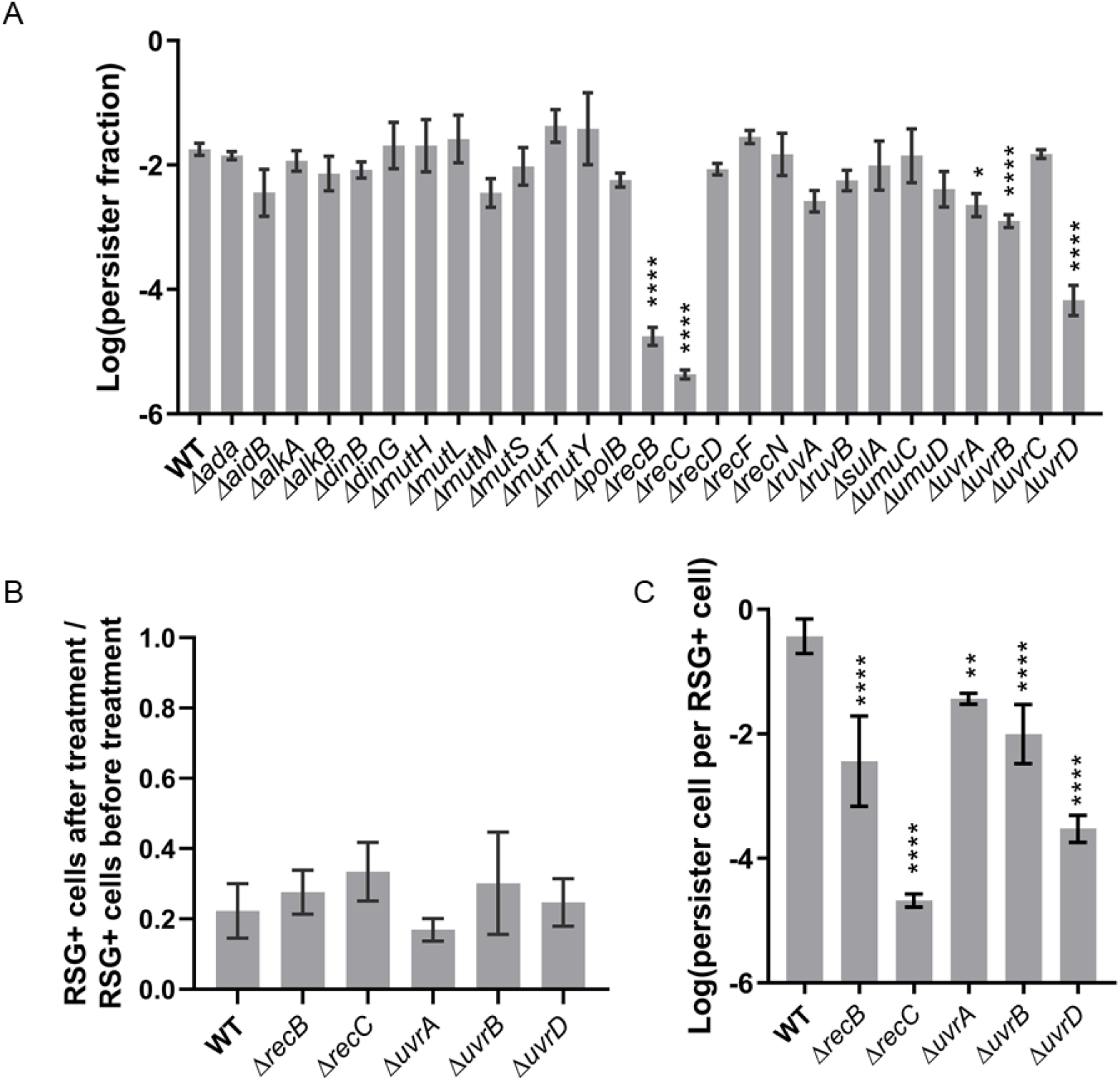
*ΔuvrB, ΔuvrD* and *ΔrecC* mutants have an impaired awakening. (A) Persister fractions after ofloxacin treatment of stationary phase cells. Values result from n ≥ 3 biological repeats and are represented as mean ± SEM. Data are log-transformed. Asterisks indicate significant difference compared to the wild type (*E. coli* BW25513) (**** P < 0.0001; ** P < 0.01). (B) The number of metabolically active cells after ofloxacin treatment over the number of metabolically active cells before ofloxacin treatment is not significant different in Δ*recB*, Δ*recC*, Δ*uvrA*, Δ*uvrB* and Δ*uvrD* compared to the wild type. Metabolic activity is measured using RSG staining. Values result from n ≥ 3 biological repeats and are represented as mean ± SEM. (C) After treatment with ofloxacin, the number of regrowing persister cells per metabolically active cell (RSG+) is significantly lower for *ΔuvrA*, Δ*uvrB*, Δ*uvrD*, Δ*recB* and Δ*recC* compared to the wild type. Values result from n ≥ 3 biological repeats and are represented as mean ± SEM. Asterisks indicate significant difference compared to the wild type (*E. coli* BW25513) (**** P < 0.0001; ** P < 0.01).

Persister fractions assessed using plate counts result from the combined effect of rates of persister state entry and exit. To distinguish whether the absence of *uvrA, uvrB* and *uvrD* impedes persister formation or awakening, we first assessed the number of metabolically active cells before treatment compared to the number of metabolically active cells after antibiotic treatment using the metabolic activity sensor redox sensor green (RSG). The green fluorescence is positively correlated with the metabolic activity of the cell. No significant difference of the ratio of RSG+ cells after and before treatment between the *ΔrecB*, Δ*recC*, Δ*uvrA*, Δ*uvrB* and Δ*uvrD* mutants and the wild type was observed (Fig 1B). This demonstrates that an equal fraction of cells is metabolically active after the ofloxacin treatment, suggesting that the deletions do not interfere with direct cell survival following antibiotic treatment. Second, the number of persister cells per metabolically active cell after antibiotic treatment was calculated, as published previously (18). For this, the RSG staining was performed in parallel with CFU counting. Figure 1C clearly shows that absence of *recB, recC, uvrA, uvrB* and *uvrD* results in significantly less persisters per metabolically active cells (RSG+) after treatment, indicative of an impaired awakening efficiency. Combined, these data demonstrate that absence of *recB, recC, uvrA, uvrB* and *uvrD* disrupts successful persister awakening in ofloxacin-treated cells. As the molecular role of *recBC* has been studied before in relation to persistence (24, 44), we further focused on the *uvr* genes.

### 2. Single-cell analysis during recovery reveals different awakening kinetics between antibiotic-sensitive and persister cells

To further confirm the role of UvrA, UvrB and UvrD in persister awakening, detailed single-cell awakening kinetics of the corresponding deletion mutants were quantified using time-lapse microscopy (movie S1-S2). In agreement with previous reports (22, 24), the majority of the cells starts elongating during recovery as a response to the ofloxacin treatment (Fig. 2A). However, only persister cells are able to divide multiple times (n > 4) and to generate viable progeny.

**Figure 2.**
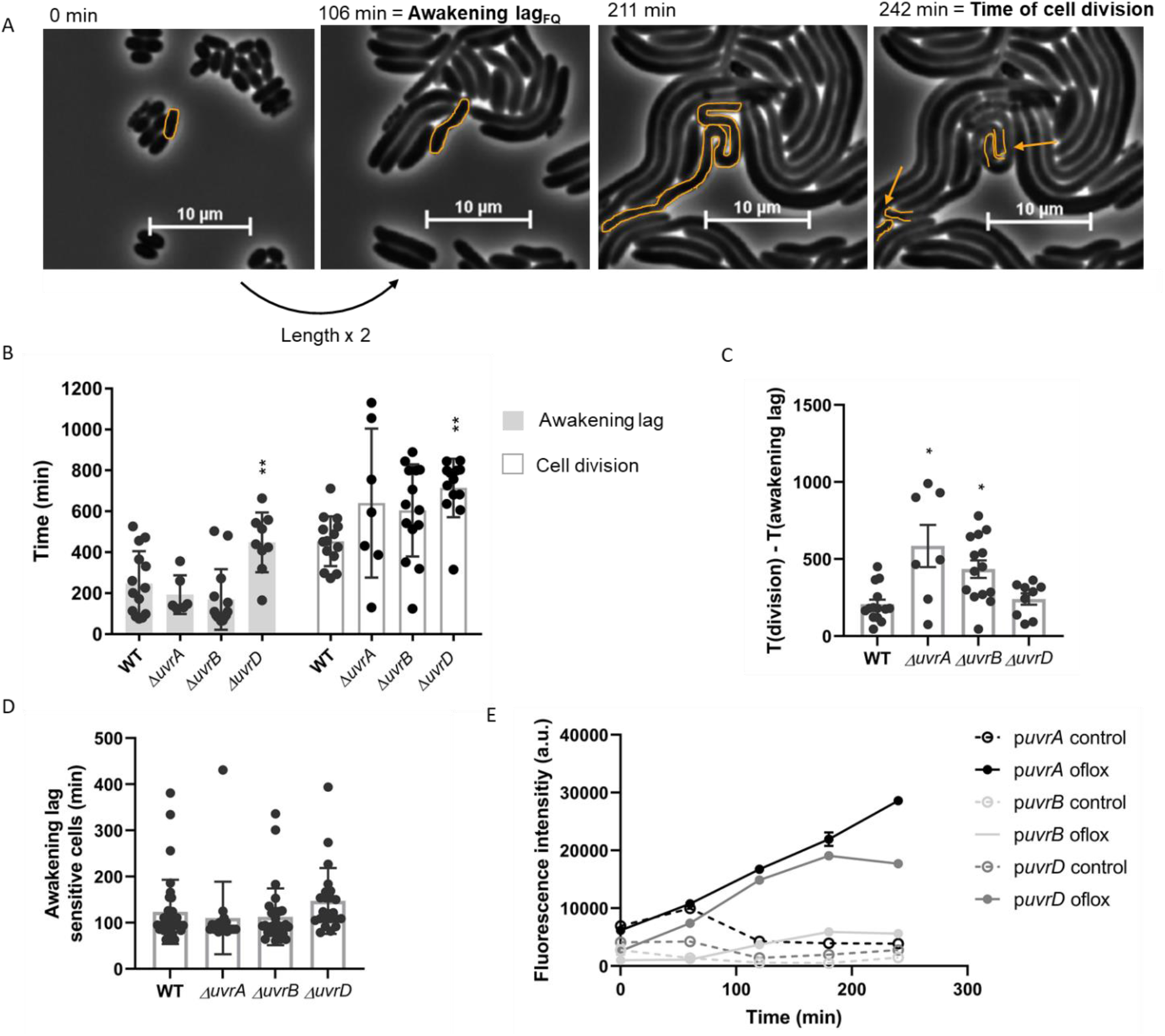
Awakening kinetics of wild-type *E. coli*, Δ*recB*, Δ*recC*, Δ*uvrA*, Δ*uvrB* and Δ*uvrD*. (A) Representative example of wild-type cells after ofloxacin treatment, growing on an agar pad with fresh nutrients. Almost all cells start to elongate after antibiotic treatment, but only persister cells undergo multiple rounds of cell divisions (> 2). The awakening lag is determined as the time point when the cell reaches twofold its initial size. The yellow arrows indicate the cell division planes. (B) Awakening lag and cell division for persister cells. Δ*uvrD* persister cells wake up significantly later and have a stalled cell division compared to wild type persister cells. Error bars are represented as mean ± SEM. Asterisks indicate significant difference (** P < 0.01). (C) The difference between the awakening lag and the time of cell division is significantly longer in absence of UvrA and UvrB. Error bars are represented as mean ± SEM. Asterisks indicate significant difference ( * P < 0.05). (D) The difference in awakening lag is specific for persister cells, as no significant difference is detected in awakening lag of the antibiotic-sensitive cells. Error bars are represented as mean ± SEM. (E) Following ofloxacin treatment, wild-type strains containing *pUvrA-gfp, pUvrB-gfp* and *pUvrD-gfp* were washed and diluted in LB, after which fluorescence intensity was assessed after 0, 60, 120, 180 and 240 minutes of incubation in fresh medium using flow cytometry. As a control, fluorescence intensities of untreated strains were taken. Values result from n ≥ 3 biological repeats and are represented as mean ± SEM.

Elongation of both persister cells and antibiotic-sensitive cells was monitored as a function of time, resulting in detailed quantitative assessment of awakening kinetics for both populations. Because persister cells strongly elongate before cell division following fluoroquinolone treatment, we propose that the total awakening time is characterized by two different phenotypic hallmarks: the awakening lag, defined as the time the cell needs to reach twice its initial size, and the time of cell division.

Single-cell awakening kinetics reveal that the awakening lag of individual persister cells is highly variable, ranging from 75 to 526 minutes in the wild type. Awakening lags of wild-type persisters are normally distributed, pointing towards underlying molecular mechanisms (45). In addition, during the awakening lag, the cellular elongation rate is higher in sensitive cells compared to persister cells (Fig. S2A, S2B and S2C). Absence of UvrA and UvrB does not interfere with the awakening lag (Fig. 2B), but significantly increases the time between the awakening lag and cell division (Fig. 2C). In contrast, absence of UvrD strongly increases the awakening lag. By measuring cell length as a function of time during the awakening lag in persister cells, it is clear that this increase in awakening lag is not caused by a slower cell elongation of the mutant, but a strong delayed initiation of cell elongation compared to the wild type (Fig. S2D). As this effect is specific for Δ*uvrD*, this might point towards a role for UvrD-mediated RNA polymerase backtracking in the awakening lag of persister cells, which occurs independently of UvrA2B (37). Absence of UvrD also increases the time of cell division in persister cells (Fig. 2B), although after this extended awakening lag, the time to cell division is comparable to that of the wild type (Fig. 2C). Importantly, this difference in the awakening lag between wild-type persister cells and *ΔuvrD* persisters is specific for persister cells, as no significant difference is observed in the antibiotic-sensitive cells (Fig. 2D).

Previous work demonstrated that expression of repair genes is important during the recovery period following ofloxacin treatment (21, 23). Therefore, expression of promoter-gene fusions of the *uvr* genes with *gfp* (46) was monitored as a function of time for the duration of the awakening lag in the wild type following ofloxacin treatment (Fig. 2E). Expression of *uvrA, uvrB* and *uvrD* is induced following ofloxacin treatment in the bulk of the population, although expression of *uvrA* and *uvrD* is more elevated. These data demonstrate that NER is expressed following ofloxacin treatment, and that absence of this repair pathway strongly influences persister awakening kinetics. In a next set of experiments, we investigated the molecular role of NER in persister awakening.

### 3. Persister cells use NER to repair oxidative DNA-damage

Fluoroquinolones such as ofloxacin result in the formation of DSB, which are repaired via HR (21, 24). The molecular role for UvrA, UvrB and UvrD in persister survival after ofloxacin treatment is therefore likely indirect, as NER does not contribute to DSB repair. NER is involved in the repair of a broad variety of other DNA lesions which are detected either directly via the UvrA2B complex, or indirectly via transcription-coupled repair (TCR) (26, 27, 29, 30, 37). One of the well-known damages directly detected and repaired by NER are pyrimidine dimers which are formed following UV radiation. These pyrimidine dimers are, however, not formed following ofloxacin treatment (Fig. S3).

Several studies demonstrate that antibiotics induce the formation of reactive oxygen species (ROS) (47–49). Accumulation of ROS induces several types of DNA lesions, which can induce cell death if left unrepaired or when repair is unsuccessful, for example by a defective NER machinery (48, 50–52). We therefore first investigated whether ROS are induced in our set-up. ROS production after ofloxacin treatment was confirmed using the non-fluorescent precursor H_2_DCFDA (2,7-dichlorodihydrofluorescein) (Fig. 3A), which can be oxidized to the green-fluorescent molecule H_2_DCF. Accumulation of ROS can result in DNA damage, for example via the incorporation of oxidized nucleotides (47, 52), with 8-oxoguanine as the major product (53). To investigate whether the observed increase in ROS following ofloxacin treatment results in oxidative DNA lesions, the formation of 8-oxoguanine was assessed after ofloxacin treatment and during recovery. Therefore, antibiotic-treated cells were fixed after 0, 1 and 3h during recovery in fresh medium, which corresponds to the awakening lag in the persister cells. Untreated cells were used as a control. Following fixation, anti-8-oxoguanine antibodies were applied and the cells were visualized under the microscope, after which the fluorescence intensity relative to the control cells was calculated (Fig. 3B). Here, an increase of 8-oxoguanine nucleotides is observed after 0, 1 and 3h recovery in fresh nutrients, confirming that ofloxacin treatment results in the accumulation of an oxidized nucleotide pool, leading to oxidative DNA damage upon incorporation.

**Figure 3.**
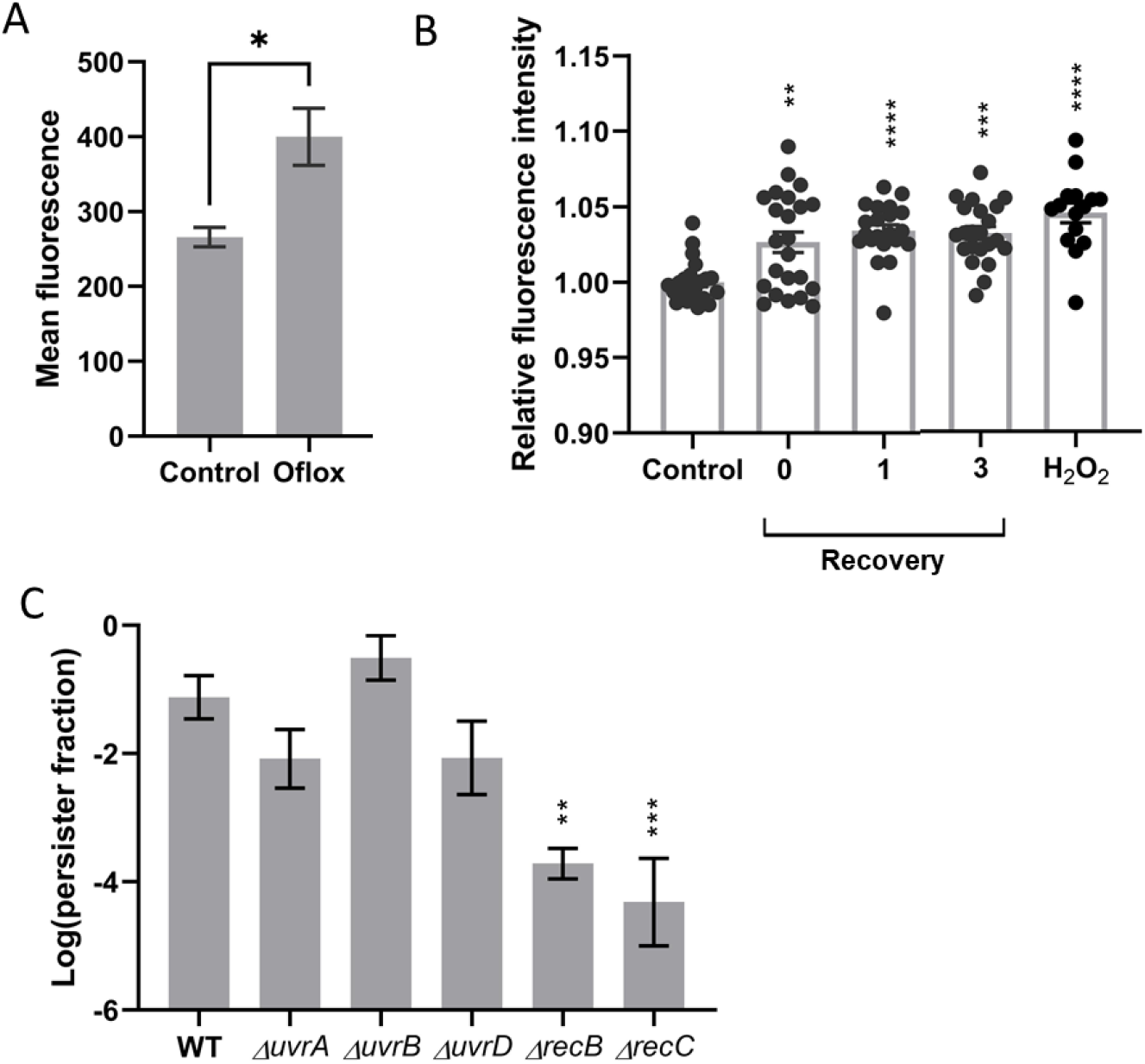
Repair of oxidative DNA damage influences persister survival. (A) ROS production after ofloxacin treatment was confirmed using the non-fluorescent precursor H_2_DCFDA, which emits green light upon oxidation to H_2_DCF. Green fluorescence was measured using flow cytometry, and the mean fluorescence of the population was calculated. Values result from n ≥ 3 biological repeats and are represented as mean ± SEM. Asterisks indicate significant difference (** P < 0.01). (B) Ofloxacin treatment results in the formation of 8-oxoguanine. Primary anti-8-oxoguanine antibodies were used, after which an Alexa Fluor secondary antibody, linked with a fluorescent protein, was used to quantify 8-oxoguanine formation. The mean fluorescence of the cell was measured using fluorescent microscopy. Values result from n ≥ 3 biological repeats and are represented as mean ± SEM. Asterisks indicate significant difference with the untreated control condition (**** P < 0.0001; *** P < 0.001; ** P < 0.01). (C) Samples were grown and treated with ofloxacin in strict anaerobic conditions. Data are log-transformed. Values result from n ≥ 3 biological repeats and are represented as mean ± SEM. Asterisks indicate significant difference compared to the wild type (*** P < 0.001; ** P < 0.01).

In a next step, we assessed the essential role of NER in the repair of oxidative DNA damage. Therefore, a persister assay was performed in anaerobic conditions to prevent the formation of ROS during ofloxacin treatment and persisters were quantified in different NER mutant backgrounds (Fig. 3C). To confirm the accumulation of DSB in anaerobic conditions, strains devoid of RecB and RecC, mediating DSB repair via HR, were taken as a control. Under anaerobic conditions, a significant decreased persister fraction is observed for Δ*recB* and Δ*recC*, but not for Δ*uvrA*, Δ*uvrB or* Δ*uvrD*, pointing towards a role for NER in oxidative DNA damage-repair during persister recovery, and confirming that NER does not have a function in DSB repair.

### 4. TCR underlies the variation in awakening lag in persister cells

In *E. coli*, different mechanisms are at play to detect and repair different types of oxidative DNA damage (27, 54). Previous experiments demonstrate that persister survival following ofloxacin treatment is based on NER-mediated repair of oxidative DNA damage. NER-mediated repair can occur directly, if the UvrA2B complex identifies the lesion, or indirectly following detection by transcription-coupled repair (TCR), where a stalled or impeded RNA polymerase is recognized and displaced by Mfd (31, 55) or moved backwards by UvrD (37). Importantly, single-cell time lapse microscopy (Fig. 2B) demonstrates that only the absence of UvrD strongly increases the awakening lag of persister cells, but not the absence of UvrA and UvrB. This implies that the role of UvrD in TCR, rather than its role in NER, governs the awakening lag. This might suggest that changes in TCR functioning will alter awakening kinetics, rather than antibiotic survival. To confirm this hypothesis, persister assays were performed on several deletion mutants influencing TCR: Δ*mfd*Δ*uvrD*, Δ*mfd* and Δ*greA*. In absence of UvrD and Mfd, the cell is devoid of a dedicated RNA polymerase displacement factor, ruling out TCR. Furthermore, GreA is an anti-backtracking factor (37, 56), of which deletion results in a backtracking-prone phenotype (37). No significant difference in survival of these mutants was observed compared to the wild type (Fig. 4A). These data demonstrate that oxidative DNA damage in persisters following ofloxacin treatment is not necessarily lethal. Of note, Δ*mfd*Δ*uvrD* has an increased persister fraction compared to Δ*uvrD*, indicating that in absence of TCR, UvrA_2_B is not able to recognize the damage. In these conditions, repair is not initiated and survival is not impaired by the absence of UvrD.

**Figure 4.**
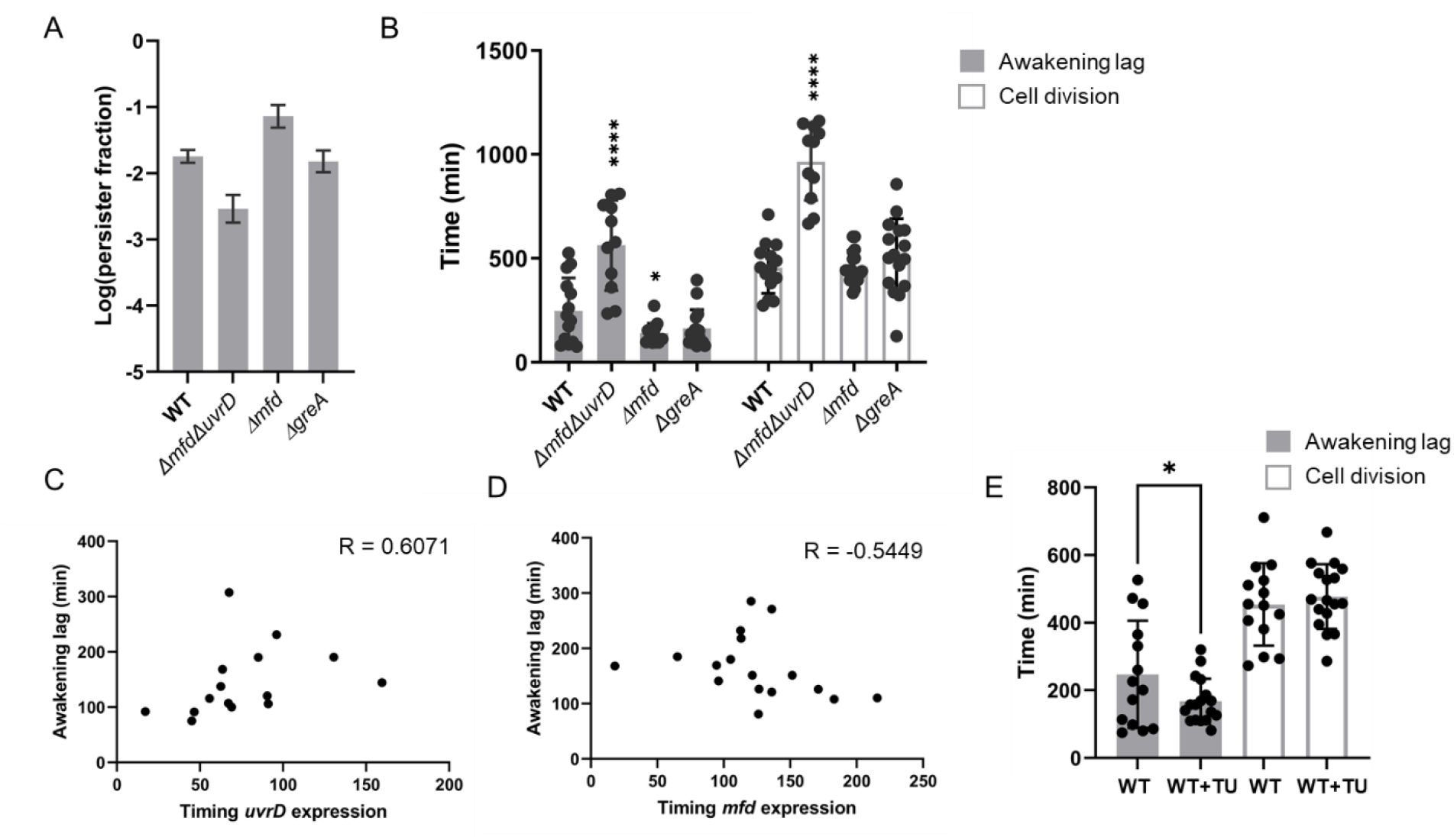
Variation in awakening lags in persister cells is caused by TCR. (A) Persister fractions after ofloxacin treatment of Δ*mfd*Δ*uvrD*, Δ*mfd* and Δ*greA*. Values result from n ≥ 3 biological repeats and are represented as mean ± SEM. Data are log-transformed. Asterisks indicate significant difference compared to the wild type (*E. coli* BW25513) (**** P < 0.0001). (B) Single-cell awakening kinetics of persister cells following recovery on LB agar pads after ofloxacin treatment. Δ*mfd*Δ*uvrD* persisters have a significantly increased awakening lag compared to wild-type persister cells. Δ*mfd* persisters have a significantly shorter awakening lag compared to wild-type persister cells. Error bars are represented as mean ± sd. Asterisks indicate statistical significance (**** P < 0.0001, * P < 0.05) compared to the wild type. A downward trend for Δ*greA* persisters is observed, but it is not significant (p = 0.099). The variation of the awakening lags of Δ*mfd*- and Δ*greA* persisters differs significantly compared to the wild type (Levene’s test, p = 0.00016 and p = 0.013, respectively). (C) Wild-type strains containing pUvrD-gfp were treated with ofloxacin, after which their recovery was followed at the single-cell level using time lapse microscopy. Fluorescence intensity and cell length were followed in time for 15 persister cells. The time of *uvrD* expression was significantly correlated with the awakening lag (p = 0.0186). Correlation was calculated using Spearman (R = 0.6071). (D) Wild-type strains containing pMfd-gfp were treated with ofloxacin, after which their recovery was followed at the single-cell level using time lapse microscopy. Fluorescence intensity and cell length were followed in time for 17 persister cells. The time of *mfd* expression was significantly correlated with the awakening lag (p = 0.0158). Correlation was calculated using Spearman (R = −0.5449). (E) Single-cell awakening kinetics of wild-type persister cells with and without TU treatment following recovery on LB agar pads after ofloxacin treatment. Cells treated with TU have a significantly shorter awakening lag compared to the untreated control. Error bars are represented as mean ± sd. Asterisks indicate statistical significance (* P < 0.05).

Next, single-cell awakening lags were determined for Δ*mfd*Δ*uvrD*, Δ*mfd* and Δ*greA* (movie S3-S4). The Δ*mfd*Δ*uvrD* mutant has a strongly increased awakening lag (Fig. 4B), which is even more increased compared to Δ*uvrD* persisters (Fig. 2A), presumably caused by the absence of a dedicated RNA displacement factor. Strikingly, Δ*mfd*-persister cells have a significantly shorter awakening lag compared to wild-type persister cells (Fig. 4B), which can be explained by the overall backtracking of the RNA polymerase in this mutant, resulting in the fast resumption of transcription. Although not significant (p = 0.099), a downward trend, similar to Δ*mfd*, is observed for Δ*greA*-persisters. This is in agreement with the backtracking-prone phenotype of Δ*greA* (37), of which the phenotype is weaker compared to complete absence of Mfd. Absence of both Mfd and GreA results in significantly less variation in awakening lag compared to wild-type cells (Levene’s test, p = 0.00016 and p = 0.013, respectively). Importantly, no significant difference in awakening lag is observed in the sensitive cells for Δ*mfd* and Δ*greA* (Fig. S4A). Interestingly, we also observe a significant difference in awakening lag in antibiotic sensitive cells when we compare wild-type and Δ*mfd*Δ*uvrD* backgrounds. However, the difference with the wild-type is strikingly higher in persister cells, demonstrating a persister-specific effect.

These data suggest that stalling of the RNA polymerase by oxidative DNA damage influences the awakening lag via an interplay between the two TCR pathways, orchestrated by UvrD and Mfd. In absence of Mfd, a stalled RNA polymerase will be rescued by UvrD-mediated backtracking. The damage will be repaired and transcription can proceed swiftly as UvrD will backtrack the RNA polymerase (37), which will result in the observed short awakening lag and the lower variation (Fig. 4B). In absence of UvrD, Mfd will displace the RNA polymerase and transcription has to restart, resulting in the observed strongly increased awakening lag (Fig. 2B). To further validate the competitive interaction between UvrD- and Mfd-mediated TCR and their influence on the awakening lag of the persister cells, expression of *uvrD* and *mfd* was monitored in persister cells during recovery using time-lapse microscopy. A positive correlation is observed between the timing of *uvrD* expression and the awakening lag at single-cell level (Fig. 4C), suggesting that early expression of *uvrD* is correlated with a short awakening lag. Furthermore, a negative correlation is observed between the timing of *mfd* expression and the awakening lag, suggesting that early *mfd* expression during persister recovery is correlated with a long awakening lag (Fig. 4D). Strikingly, persister cells have an increased expression of *mfd*, but not *uvrD*, following ofloxacin treatment compared to antibiotic-sensitive cells (Fig. S4B, S4C).

Previous data demonstrate that TCR alters the awakening lag of the persister cell in response to oxidative DNA damage. To further corroborate the influence of oxidative DNA damage on the awakening lag of persister cells, ofloxacin treatment and recovery of wild-type cells were performed in combination with the hydroxyl radical scavenger thiourea (TU) (57). Hydroxyl radicals are the main contributor to the formation of 8-oxoguanine. Treatment with TU results in a significantly decreased awakening lag (Fig. 4E), but again, did not alter persister survival (Fig. S4D). Consistent with previous data, a significantly decreased variation in awakening lag is observed in wild-type persister cells treated with TU (Levene’s test, p = 0.0047).

Combined, these data demonstrate that oxidative DNA damage does not necessarily cause cell death in a wild-type background, but initiates TCR-mediated NER, resulting in heterogeneity of the awakening lag in persister cells. If repair fails, for example when the NER machinery is defective, survival is impaired.

### 5. Mfd-mediated TCR plays a role in the emergence of resistant mutants

If left unrepaired, oxidative DNA damage such as 8-oxoguanine can be mutagenic (52). Previous work has established that persisters form a pool from which resistant mutants emerge (7–9). Furthermore, Mfd is an evolvability factor that increases mutation rates and has been linked with the emergence of antibiotic resistance (35, 36). Therefore, to address the role of oxidative stress and oxidative DNA damage repair on the accumulation of mutations in persister cells, a fluctuation assay was performed (58). Here, the distribution of the emergence of mutants in multiple parallel cultures is determined, after which the mutation rate is inferred from this distribution of mutants (58, 59). The mutation rates in wild-type, Δ*mfd*- or Δ*uvrD*-backgrounds did not differ between untreated and treated populations (Fig. 5A), meaning that the overall probability of a mutation occurring per cell division is not different between untreated and ofloxacin-treated cells (59, 60). However, an increased mutation rate is observed in Δ*uvrD*, both in untreated and treated cultures, compared to the wild-type. An increased mutation rate in Δ*uvrD* was observed before (61) and was shown to be the result of the role of UvrD in methyl-directed mismatch repair (MMR), which corrects replication errors (40, 62).

**Fig. 5.**
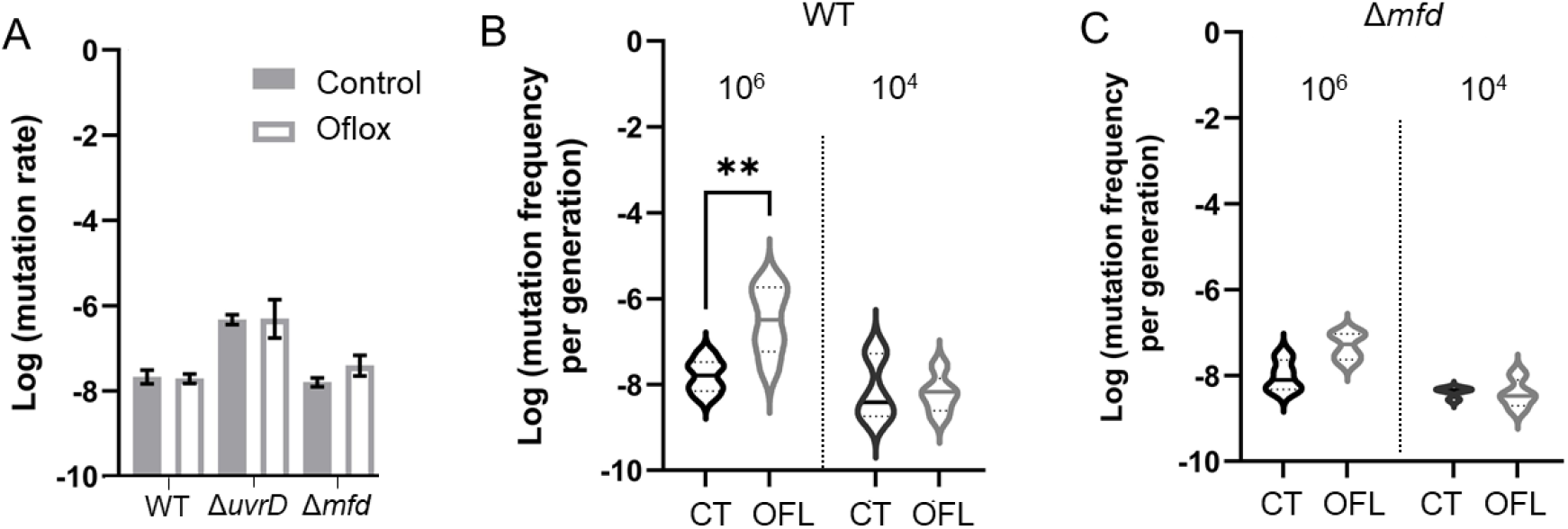
The persister population contains different subpopulations with different mutation rates. (A) No significant difference is observed in mutation rates, inferred by performing a fluctuation assay, between untreated (control) and ofloxacin-treated populations. Values result from n ≥ 3 biological repeats and are represented as mean ± SEM. (B) Taking an increased bottleneck population (10^6^) following ofloxacin survival significantly increases the mutation frequency per generation in the wild type. Values result from n ≥ 3 biological repeats and are represented as mean ± SEM. (C) Taking an increased bottleneck population (10^6^) following ofloxacin survival significantly does not increase the mutation frequency per generation in absence of Mfd. Values result from n ≥ 3 biological repeats and are represented as mean ± SEM.

Importantly, a previous report demonstrated that a population treated with fluoroquinolones has a higher mutation frequency (63). The mutation frequency is determined as the proportion of mutants in a culture and is strongly biased if a mutational event occurs early in exponential phase (59). We hypothesized that this discrepancy was the result of a higher mutation rate occurring sporadically in a very limited number of persister cells. Therefore, different bottlenecks of populations were tested. Populations were treated with antibiotics or left untreated, and subsequently 10^4^ or 10^6^ cells were transferred to fresh medium. The mutation frequency was assessed after overnight incubation (Fig. 5B). No significant increase in mutation frequency is found upon recovering 10^4^ cells, which is the same population size as that used in the fluctuation assay in Fig. 5A. However, when 10^6^ cells are taken, an increased number of mutants per generation upon ofloxacin treatment in the wild type is observed compared to the untreated control (Fig. 5B). Given the promutagenic role of Mfd and the fact that absence of Mfd decreases the formation of resistant mutants (35), we hypothesized that Mfd might be responsible for these sporadic mutational bursts in persister cells. In absence of Mfd, no significant difference in the mutation frequency per population bottleneck is observed (p = 0.22), indicating that Mfd is indeed responsible for bursts in mutational events in persister cells following ofloxacin treatment (Fig. 5C).

According to these data, we can infer that most persisters do not have an altered mutation rate. However, a small subpopulation of persister cells have an elevated mutation rate and are more likely to form resistant mutants, triggered by Mfd.

## Discussion

To date, persistence research has mainly focused on mechanisms involved in the induction of the persister state. From a medical point of view, targeting persister awakening is much more interesting than targeting persister formation as a bacterial infection likely already accommodates persister cells at the moment of treatment. As persisters accumulate damage during fluoroquinolone treatment, the ensuing recovery response is a promising target for an anti-persister therapy. Therefore, detailed insights in the molecular mechanisms underlying damage repair during persister awakening are of critical importance.

In this work, we deepened the role of DNA damage repair in persister survival after ofloxacin treatment. Fluoroquinolones such as ofloxacin result in the formation of DSB by blocking DNA-gyrase functioning. The formation of these DSB in persister cells was reported previously (21) and it was demonstrated that persister cells containing two chromosomes are more likely to survive fluoroquinolone treatment, resulting from the ability to perform HR (24). This is in agreement with our observation that the presence of RecB and RecC is important for successful persister survival, as in *E. coli* HR is predominantly orchestrated via the RecBCD pathway (64).

In addition to HR, our results clearly point towards a prominent role for NER in persister survival by oxidative DNA damage repair. Our work and others have demonstrated that antibiotics induce the formation of ROS (47, 48, 50, 51). Whether or not this accumulation results in cell death is under debate (49, 65). Our results demonstrate that the oxidative DNA damage following ofloxacin treatment does not induce cell death directly in a wild-type background, but can cause cell death if repair is corrupted, which is in line with previous reports (52, 66). Although the repair of oxidative DNA damage is often attributed to base-excision repair mechanisms (BER) (52), recent work has demonstrated that NER is important for survival of oxidative stress when the level of damage is high (27). Our results demonstrate that in *E. coli* persisters, the NER pathway repairs oxidative DNA damage, as absence of UvrA, UvrB and UvrD significantly lowers persister survival during recovery in aerobic conditions. Of note, UvrD was linked with decreased antibiotic survival in *E. coli* MG1655 (23, 44), suggesting a general mechanism.

Our results show that detection of the oxidative DNA damage occurs via TCR. Although under debate (29, 55), oxidative DNA damage can stall or pause RNA polymerase, which is recognized and displaced by Mfd (31, 55) or moved backwards by UvrD (37) to expose the site of the lesion and enable repair. In both cases, this repair is orchestrated via NER. While transcription can swiftly proceed after repair if the RNA polymerase is moved backwards by UvrD, transcription terminates following Mfd-mediated forward movement of the RNA polymerase (33, 37, 56). Hence, the molecular mechanisms underlying movement of the RNA polymerase dramatically change transcription dynamics, and as a result, the awakening lag of the persister cells. UvrD-mediated backtracking results in a short awakening lag, while Mfd-mediated displacement of the RNA polymerase results in a long awakening lag of the persister cells. Importantly, the role of UvrD is twofold: while our data demonstrate that its role in NER-mediated repair as a helicase will influence persister survival, its role in TCR influences the awakening lag. At the population level, these differences in transcription dynamics and competition between these systems will result in heterogeneous persister awakening, as was observed before (22, 24). We can speculate that this phenotypic variability aids in survival of the persister population upon antibiotic encounter, as it diverges the timing that persister cells become antibiotic-sensitive again (67).

Some of the oxidative DNA damage, such as 8-oxoguanine, is mutagenic upon incorporation in the DNA. Previous work has suggested that the accumulation of ROS following antibiotic treatment plays a role in resistance development (9, 68). In addition, increased oxidative stress in biofilms increases mutation frequencies in *Staphylococcus aureus* (69). Our results demonstrate that following antibiotic treatment, the majority of the persister cells repair the oxidative DNA damage, and have a mutation rate equal to the untreated population. However, a small subpopulation of persister cells has a higher mutation rate, triggered by Mfd, resulting in the emergence of resistant mutants (Figure 6). Indeed, in addition to its role in repairing DNA damage, Mfd was found to play a role in the evolution of antimicrobial resistance in *Salmonella typhimurium, Bacillus subtilis, M. tuberculosis* and *Campylobacter jejuni* (35, 70) and in stationary-phase mutagenesis in *Pseudomonas putida* and *Bacillus subtilis* (61, 71). This promutagenic function presumably works via the formation of R-loops, inherent to transcription, where one strand of DNA is annealed to a complementary RNA strand instead of its complementary DNA strand (36). These R-loops are precursors for both DSB and double-stranded ends (DSE), which are mutagenic both during replication and in starvation-stressed cells (72). In stationary phase, TCR predominantly occurs via UvrD-mediated backtracking (37, 38). Hence, these jackpot events are rare under these conditions, however, they have a high biological relevance if the number of residing persisters at an infection site following antibiotic treatment is high. Importantly, subpopulations with different mutation rates were observed before and it was postulated to be a bet-hedging strategy (67, 68, 73). Our work links extended awakening lags with a higher mutation rate (Figure 6), shifting the balance between genome integrity and plasticity in time, which might be a strategy of the persister population to preserve genome integrity of the population if mutagenesis is not needed (74).

**Figure 6.**
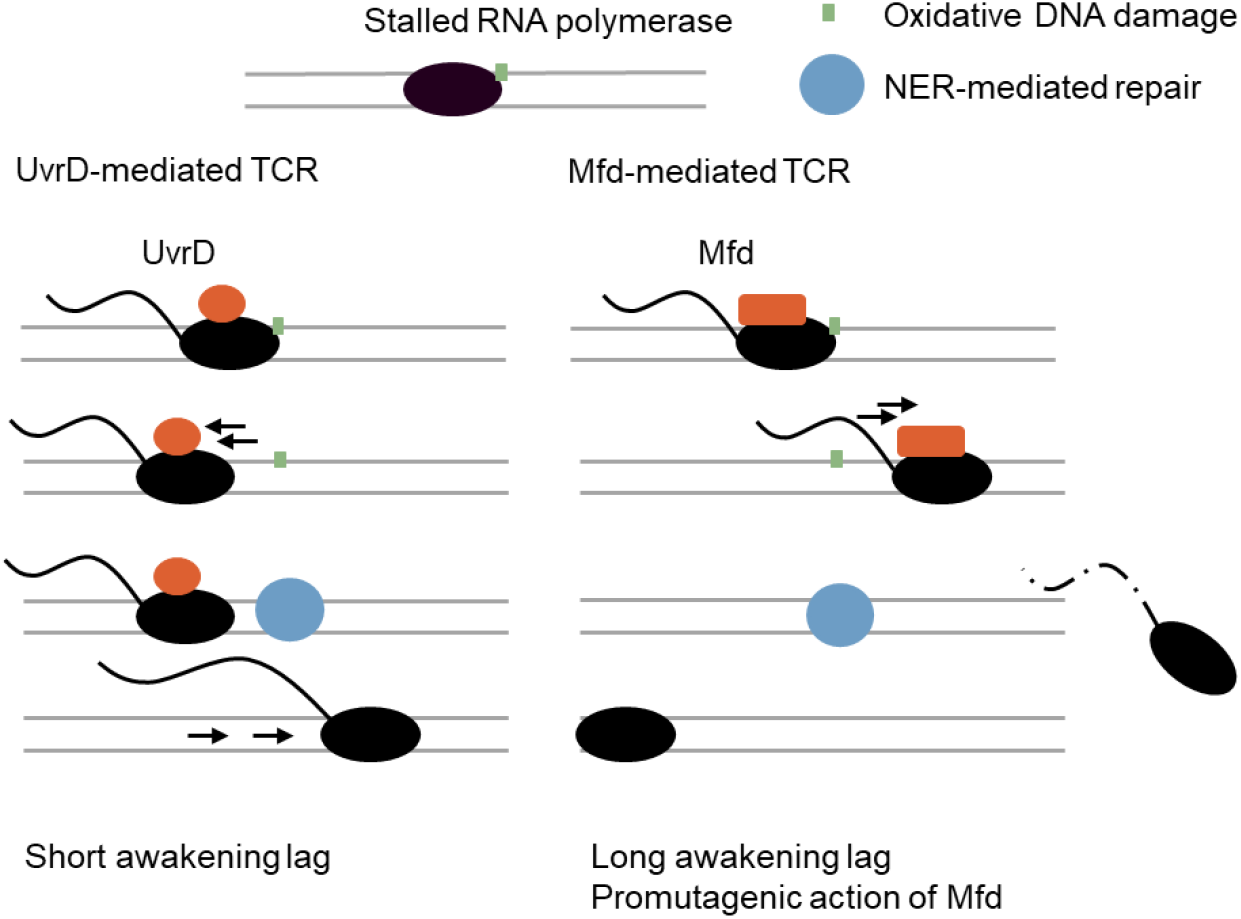
Model of the influence of TCR on the awakening lag of persister cells following fluoroquinolone treatment. Upon stalling of the RNA polymerase by oxidative DNA damage, TCR is initiated to remove the RNA polymerase from the site of the lesion (green square), enabling NER to repair the damage. UvrD-mediated movement of the RNA polymerase results in direct proceeding of transcription, resulting in a short awakening lag of the persister cell. Mfd-mediated displacement of the RNA polymerase terminates transcription, increasing the awakening lag of the persister cell. In addition, the promutagenic factor Mfd will result in jackpot events in a small fraction of the persister cells, leading to antibiotic resistance.

Persister cells and sensitive cells were found to have three major behavioral differences following antibiotic treatment. First, persister cells have a significantly decreased rate of elongation compared to antibiotic-sensitive cells. Second, our data demonstrate that the influence of TCR on the awakening lag is specific for persister cells. However, the observation that sensitive cells also elongate strongly suggests that transcription is initiated as well. Third, persister cells have a significantly increased *mfd* expression compared to sensitive cells immediately after ofloxacin treatment. In another study, increased expression of *mfd* was also observed following ciprofloxacin treatment in *C. jejuni* (70). Presumably, these differences will change awakening kinetics and alter transcription dynamics, resulting in the observed difference between persister cells and sensitive cells. It is tempting to speculate that alteration of transcription speed increases the chance of DSB repair by halting growth resumption, thereby increasing the chance of survival (23, 75).

The findings reported here shed new light on the role of NER in successful persister recovery following fluoroquinolone treatment and demonstrate that TCR dictates the heterogeneity of awakening lag of persister cells. In addition, we reveal the presence of a subpopulation of persisters with a higher mutation rate, driven by the promutagenic factor Mfd (Figure 6). Finally, an increased awakening lag seems correlated with a higher mutation rate. This way, the chance of survival of the bacterial population is increased, while genome integrity is maintained.

## Materials and methods

### Bacterial strains, growth conditions and plasmids

The *E. coli* strain BW25113 was used for all experiments (76). Bacteria were grown in lysogeny broth (LB) at 37°C under orbital shaking (200 rpm). When needed, kanamycin was added (40 μg mL^-1^ for genomic deletion mutants and 20 μg mL^-1^ for the promoter fusions) to the culture.

Construction of pUvrB was performed by PCR amplification of the intergenic region (500 bp) preceding the start codon of the *uvrB* gene using primers **ACT GCT CGA GCC ATT CTG TAT TTG GTT AAA and CAC CGG ATC CGA GTC GCT ACC TGA AGG AGT**. Construction of pRecC was performed by PCR amplification of the promoter region of *recC* using primers ACT GCT CGA GAT GTC AGC TTC CCT GAA GAA and CAC CGG ATC CAG CGG CTC CTG ACT ACT GAC. The corresponding PCR produces were digested with XhoI and BamHI and cloned in PUA66 (46). pRecB was constructed using Gibson assembly. The promoter region of *recB* was amplified using TTC TTA AAT CTA GAG GAT CCT CAC GGA CCT CAT AAT CAA CTT AAT TTT CTGT and CTT TCG TCT TCA CCT CGA GAG TGC TGT ATA AAA ATT GCG CAA TCT. The PUA66 plasmid was linearized using XhoI and BamHI. Gibson assembly was performed using the Gibson assembly master mix^®^ purchased from New England Biolabs. pUvrA and pUvrD were purified from the library constructed in (46) and transformed to BW25113.

### Persistence assay

Cells were inoculated in 5 mL LB medium and incubated for 24 h at 37°C under orbital shaking. This preculture was diluted 100-fold in 5 mL LB medium and again incubated for 16 h. Then, 990 μL of the culture was treated with 10 μL ofloxacin with a final concentration of 5 μg ml^-1^ or with 10 μL moxifloxacin with a final concentration of 2.4 μg ml^-1^. As a control, 990 μL of the culture was treated with 10 μL sterile water. Cultures were treated for 5 h under shaking conditions at 37°C. Scavenging of the hydroxyl radicals was performed by adding 150 μM thiourea during the ofloxacin treatment. The antibiotic treated samples were washed two fold in 10 mM MgSO_4_ to remove the antibiotics, after which all samples were serial diluted in 10 mM MgSO_4_ and plated out on LB agar plates. The number of colony forming units (CFU) was determined after 48 h of incubation at 37°C. The persister fraction is determined as the log_10_ transformation of the number of cells after treatment over the number of cells before treatment.

The persister assay in anaerobic conditions was performed in a Whitley DG250 anaerobic workstation (flushed with 80% N_2_, 10% CO_2_ and 10% H_2_). Prior to usage, all media were incubated for 24 h in the anaerobic workstation. Statistical comparisons were based on a one-way ANOVA compared to the wild type with a Dunnett’s *post hoc* test for multiple comparison.

### Determination of persister cells in the metabolically active fraction

To determine the number of persister cells in the metabolically active fraction, a persistence assay was performed as described above. Parallel with plating out the samples, the samples were treated with redox sensor green, purchased from Invitrogen (USA). Therefore, the control cultures were diluted 10-fold and 2 μL of this dilution was dissolved in 198 μL PBS containing RSG, according to the manufacturer’s specifications. 2 μL of the antibiotic-treated samples were directly diluted in 198 μL PBS with RSG. After 10 min incubation at room termperature, the number of metabolically active cells per mL were counted using a cytoFLEX S (Beckman Coulter, USA). Data were analyzed using CytExpert (Beckman Coulter, USA). The number of metabolically active cells after treatment was divided over the number of metabolically active cells before treatment. In addition, the number of regrowing cells on plate was divided over the number of metabolically active cells measured using the cytoFLEX S, and the data were log-transformed. Data were analyzed using a one-way ANOVA test compared to the wild type with a Dunnett’s *post hoc* analysis for multiple comparison.

### Determination of gene expression during recovery after antibiotic treatment using flow cytometry

Cells were incubated and treated with antibiotics as described for the persistence assay. After ofloxacin treatment, 500 μL of the antibiotic-treated samples were washed by centrifuging at 6000 rpm for 5 min, after which the pellet was diluted in 500 μL PBS. 10 μL of this resuspension was pipetted in an Eppendorf tube containing 990 μL LB and 20 μg mL^-1^ kanamycin. 10 μL of the control culture was diluted directly in an Eppendorf tube containing 990 μL LB and 20 μg mL^-1^ kanamycin. Samples were taken at dedicated time points and green fluorescence (488 nm) was measured in a CytoFLEX S benchtop flow cytometer. Data were analyzed using CytExpert, determining the mean green fluorescence intensity of the bacterial population.

### Single-cell analysis of persister cells and antibiotic-sensitive cells using microscopy

Cells were incubated and treated with antibiotics as described for the persistence assay. After ofloxacin (5 μg mL^-1^) treatment, cells were washed twice by centrifuging at 6000 rpm for 5 min and resuspending the cells in preheated (37°C) LB medium. After washing, 2 μL of the culture was spotted on LB agar pads. For the treatment with thiourea, 150 μM was added to 990 μL culture in combination with the ofloxacin treatment, as described before. After 5 h incubation, cells were washed twice, as described before. 150 μM thiourea was added to the agar pad. Phase contrast images and, if needed, fluorescence images were taken every 15 min using a Nikon Eclipse Ti-E inverted microscope with a CFI Plan Apochromat 100X objective with a numerical aperture of 1.45. Green fluorescent images were taken using an exposure time of 20 ms and an analog gain of 6.4. Images were analyzed using the NIS Elements Analysis 4.3 software (Nikon, Japan). The cell length was followed in time. When the cell reaches twofold its initial size, this time point was set as the awakening lag. When a cell division plane was visible, this time point was set as the time of cell division.

Normal distribution of the data was assessed using the Shapiro-Wilk test. If the data were not normally distributed, Box Cox transformation was applied to normalize the data prior to statistical testing. For Fig. 2B, data were analyzed using a one-way ANOVA test compared to the wild type with a Dunnett’s *post hoc* analysis for multiple comparison. For Fig. 2C, data were analyzed using a Kruskal-Wallis test compared to the wild type with a Dunnett’s *post hoc* analysis for multiple comparison. For Fig. 2D, data were analyzed using a one-way ANOVA test compared to the wild type with a Dunnett’s *post hoc* analysis for multiple comparison, following Box-Cox transformation (λ = −0.6) of the awakening lag to normalize the data. For the awakening lag in Fig. 4B, statistical significance was calculated by normalizing the awakening lags using Box Cox (λ = −0.1). Data were analyzed using a one-way ANOVA. For Fig. 4E, statistical significance was calculated by using a one-sided Welch’s t-test. Differences in variation were assessed using a Levene’s test. For Fig. S4A, statistical significance was calculated by normalizing the awakening lags using Box Cox (λ = −0.8). Data were analyzed using a one-way ANOVA compared to the wild type with a Dunnett’s *post hoc* analysis for multiple comparison.

An exponential curve (y=y0e^kx^) was plotted on the cell length in time for the awakening lag using Graphpad Prism, inferring the initial cellular elongation rate (k). Statistical significance was assessed using an unpaired two-sided t-test.

The fluorescence at the start of the experiment was set as the initial fluorescence. Statistical significance was assessed using a unpaired two-sided t-test. The timing of fluorescent expression was determined by plotting a one-phase decay (Y=(Y0 - Plateau)*exp(-K*X) + Plateau) on the fluorescence intensity over time (77), with a 95% confidence interval. K was constrained at 0.01264, based on the average K of 10 randomly selected bleaching curves. If the fluorescence intensity at a given time point exceeded the 95% confidence interval, this time point was set as the time point of expression. The timing of expression was correlated with the awakening lag of the persister cells using Spearman.

### Luria Delbruck fluctuation assays

Cells were incubated and treated with antibiotics as described for the persistence assay. Next, an inoculum mixture was prepared in LB containing 10^4^ cells. This inoculum mixture was plated out to verify the initial cell number. For every strain, 16 wells of a 96-well plate were filled with 200 μL of the inoculum mixture. Next, the strains were incubated for 24 h at 37°C and 200 rpm. After overnight growth, 4 of the 16 wells were diluted in MgSO_4_ and plated out on LB plates to quantify the number of cells in the culture. The content of the 12 other wells was plated on rifampicin plates (100 μg mL^-1^). Rifampicin plates were stored and incubated dark conditions. All plates were incubated at 37°C and counted after 2 days.

The total cell number (N_cells_) was calculated by taking the average of CFU mL^-1^ of the 4 non-selective plates. Mutation rates were calculated using bzrates (http://www.lcqb.upmc.fr/bzrates) and log_10_-transformed. Statistical analysis was performed using an ANOVA with Šidák correction, where the treated population was compared to the untreated population.

### Population mutation rates

Cells were incubated and treated with antibiotics as described for the persistence assay. Next, 10^6^ or 10^4^ cells were transferred to tubes containing 5 mL LB. The original culture was plated out to assess initial cell number. After 24 h incubation, a tenfold dilution series was made in MgSO_4_ and the culture was plated out on an LB plate, assessing the total number of cells. 200 μL of the culture was plated out on rifampicin plates (100 μg mL^-1^). Rifampicin plates were stored and incubated in dark conditions. All plates were incubated at 37°C and counted after two days. The number of mutants was divided over the total number of cells. To compensate for the difference in the number of cell divisions that can occur between the different population sizes, the mutation rate of the cultures was multiplied by log(100)/log(2) for the 10^6^ population. Data were analyzed using ANOVA with Šidák correction, where the treated population was compared to the untreated population.

### Measuring formation of thymine dimers

A persistence assay was performed as described before. Samples were taken immediately after antibiotic treatment, and a non-treated sample was taken as a control. To assess the influence of a recovery phase on the formation of thymine dimers, 800 μL of an untreated and antibiotic-treated sample were taken and centrifuged for 5 min at 6000 rpm, after which the pellet was dissolved in 800 μL fresh LB medium. This was done twice to assure the complete removal of ofloxacin. Samples were incubated for 30 min at 37 °C, after which they were fixated. As a positive control, 1mL of stationary phase cells were pipetted in an empty petri dish and spread out. The petri dish was kept for 30 seconds on a transilluminator, after which the cells were collected in an Eppendorf tube.

Fixation and staining of the cells was performed as described before (78), with some slight adaptations. 800 μL of the samples were centrifuged 6000 x g for 5 min, after which the pellet was dissolved ice-cold 3:1 methanol-acetic acid mixture and stored at −20°C for 5 min. After fixation, cells were centrifuged at 12000 x g for 5 min, and the pellet was washed with 70 % ethanol and stored at 4° for 45 min. Cells were centrifuged at 12000 x g for 5 min, after which the pellet was dissolved in 70% ethanol containing 0.1 M NaOH. Following centrifugation (12000 x g, 5 min), the pellet was washed twice with PBST (PBS containing 0.05 % Tween 20).

The cells were incubated with anti-thymine dimer antibody (diluted 1:2000) (RRID:AB_261597), purchased from Sigma, for 18 h at room temperature in PBST containing 1 % bovine serum albumin with orbital shaking (80 rpm). After overnight incubation, cells were washed twice with PBST. Cells were incubated with Alexa Fluor^®^ 488 goat anti-mouse IgG1 antibodies (diluted 1:10000) (RRID:AB_2535764) for 3 h at room temperature in PBST under orbital shaking (80 rpm). Secondary antibodies were purchased from ThermoFisher. Localization of the fluorescence was determined using a Nikon Eclipse Ti-E inverted microscope with a CFI Plan Apochromat 100X objective with a numerical aperture of 1.45 by spotting 2 μL of the culture on agarose agar pads (2 %). This confirmed that upon UV exposure, fluorescent foci were formed in the cell. Green fluorescent images were taken using an exposure time of 20 ms and an analog gain of 6.4. Images were analyzed using the NIS Elements Analysis 4.3 software (Nikon) by measuring the fluorescence intensity of the whole cell. For assessing the fluorescence of the population, cells were diluted 100-fold in PBS and green fluorescence (488 nm) and were measured using a cytoFLEX S (Beckman Coulter, USA). Data were analyzed using CytExpert (Beckman Coulter, USA), determining the mean green fluorescence intensity of the bacterial population. Statistical comparisons were based on ANOVA, compared to the untreated bacterial culture, with a Dunnett’s *post hoc* test for multiple comparison.

### Measuring formation of 8-oxoguanine

A persistence assay was performed as described earlier. Immediately after antibiotic treatment, 400 μL of the culture was centrifuged (6000 rpm, 5 min) and washed with fresh LB medium. Next, the culture was diluted 100-fold in an Eppendorf tube containing 1200 μL LB. Samples (200 μL) were taken immediately, or after 1, 2 and 3h incubation at 37°C.

Fixation and staining of the cells was performed as described for the detection of thymine dimers, with some optimization for the used antibodies. Ice-cold 3:1 methanol-acetic acid mixture (200 μL) was added to the samples, after which the samples were incubated on ice for 5 min. After this fixation step, samples were centrifuged at 12000 x g for 5 min, after which the pellet was dissolved in 70 % ethanol and stored at 4°C for 10 min. Cells were centrifuged at 12000 x g for 5 min, after which the pellet was dissolved in 70% ethanol containing 0.1 M NaOH. Following centrifugation (12000 x g, 5 min), the pellet was washed twice with PBST (PBS containing 0.05 % Tween 20).

The cells were incubated with anti-oxoguanine antibodies (diluted 1:2000) (RRID:AB_11210698), for 16 h at room temperature in PBST containing 2 % bovine serum albumin under orbital shaking (80 rpm). Following incubation, samples were washed twice with PBST, after which they were incubated for 2 h with Alexa Fluor^®^ 488 goat anti-mouse IgM antibodies (1:10000) (RRID:AB_2801490) dissolved in PBST containing 2 % bovine serum albumin. After incubation, samples were washed and resuspended in PBST. Cells were visualized using a Nikon Eclipse Ti-E inverted microscope with a CFI Plan Apochromat 100X objective with a numerical aperture of 1.45 by spotting 2 μL of the culture on agarose agar pads (2 %). Fluorescence was distributed over the cell, as expected. Green fluorescent images were taken using an exposure time of 20 ms and an analog gain of 6.4. Images were analyzed using the NIS Elements Analysis 4.3 software (Nikon) by measuring the mean fluorescence intensity of the cell.

Fluorescence intensities between biological repeats varied resulting from small unavoidable changes in the protocol, such as room temperature. Therefore, for every biological repeat, fluorescence intensities were divided over the average fluorescence intensity of the control sample. Data were analyzed using a one-way ANOVA test compared to the control condition (no ofloxacin treatment) with a Dunnett’s *post hoc* analysis for multiple comparison.

### Measuring formation of ROS

A persistence assay was performed as described above. 500 μL of each culture was washed and diluted in PBS containing 10 μM H_2_DCFDA (Sigma-Aldrich). Tubes were incubated at 30 min in the dark at room temperature. Fluorescence was measured using a cytoFLEX S (Beckman Coulter, USA). Data were analyzed using CytExpert (Beckman Coulter, USA), determining the mean green fluorescence intensity of the bacterial population. Statistical analysis was performed using a two-sided t-test.

## Author contributions

Conceptualization: D.W. and J.M., Methodology: D.W., Validation: D.W. Investigation: D.W., C.F. and P.M., Writing – original draft: D.W., Writing – review & editing: D.W. and J.M., Visualization: D.W., Supervision: J.M.

## Acknowledgements

We thank Prof. Chris Michiels for the use of the anaerobic workstation. The work was supported by grants from the Research Foundation Flanders (FWO; G0B0420N, G055517N), KU Leuven (C16/17/006) and VIB. D.W. received a postdoctoral mandate from the FWO (1204621N).

## Declaration of interest

The authors declare no competing interests.

## Notes

### Competing Interest Statement

The authors have declared no competing interest.

