## Supplemental information for "Transcription-coupled DNA repair underlies variation in persister awakening and the emergence of resistance"

### Supplementary figures

**Table S1:** List of genes involved in DNA damage repair

| **Gene** | **Protein** |
| --- | --- |
| *ada* | Repair of alkylated dna |
| *aidB* | Repair of alkylated dna |
| *alkA* | Repair of alkylated dna |
| *alkB* | Repair of alkylated dna |
| *dinB* | DNA polymerase IV |
| *dinG* | Helicase |
| *mutH* | Mismatch repair |
| *mutL* | Mismatch repair |
| *mutM* | Base excision repair |
| *mutS* | Mismatch repair |
| *mutY* | Base excision repair |
| *polB* | DNA polymerase II |
| *recB* | Helicase |
| *recC* | Helicase (recBCD) |
| *recD* | Helicase |
| *recF* | ssDNA binding |
| *recN* | Recombinational repair |
| *ruvA* | Holiday junction branch migration |
| *ruvB* | Holiday junction branch migration |
| *sulA* | Inhibitor of cell division |
| *umuC* | DNA polymerase V |
| *umuD* | DNA polymerase V |
| *uvrA* | Nucleotide excision repair |
| *uvrB* | Nucleotide excision repair |
| *uvrC* | Nucleotide excision repair |
| *uvrD* | Helicase |

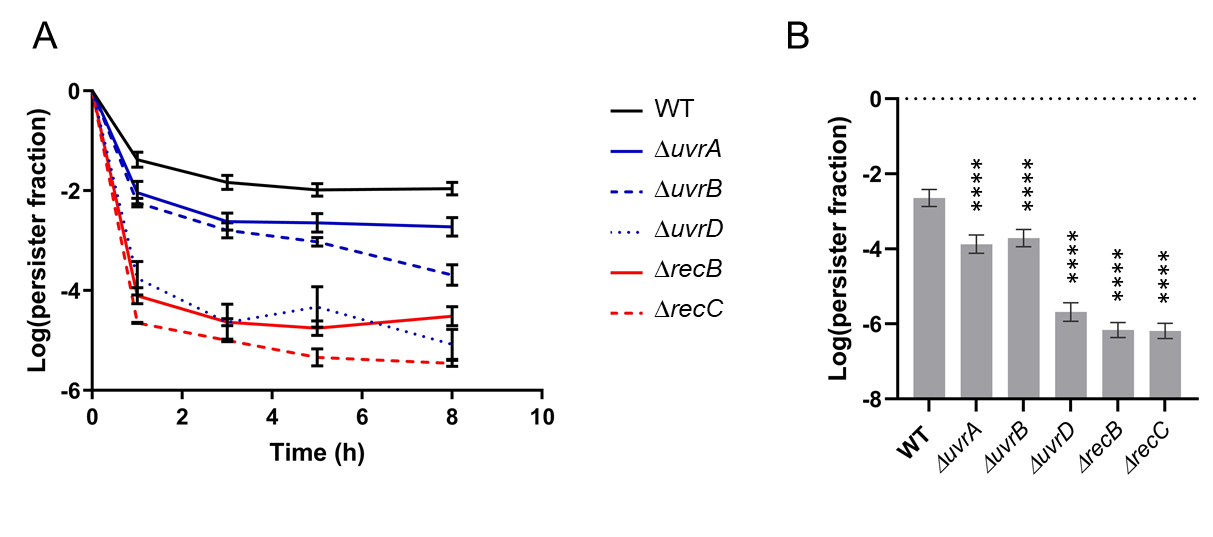

Figure S1. Absence of *uvrA, uvrB, uvrD, recB* and *recC* decreases the persister fraction. Related to Figure 1. (A) Time-kill curves were performed on *ΔuvrA*, *ΔuvrB, ΔuvrD, ΔrecB* and *ΔrecC*, demonstrating a biphasic killing curve following fluoroquinolone treatment, resulting from the presence of persister cells. Values result from n ≥ 3 biological repeats and are represented as mean ± SEM. (B) Moxifloxacin treatment of stationary phase cultures. Values result from n ≥ 3 biological repeats and are represented as mean ± SEM. Data are log-transformed. Asterisks indicate significant difference compared to the wild type (*E. coli* BW25513) (**** P < 0.0001).

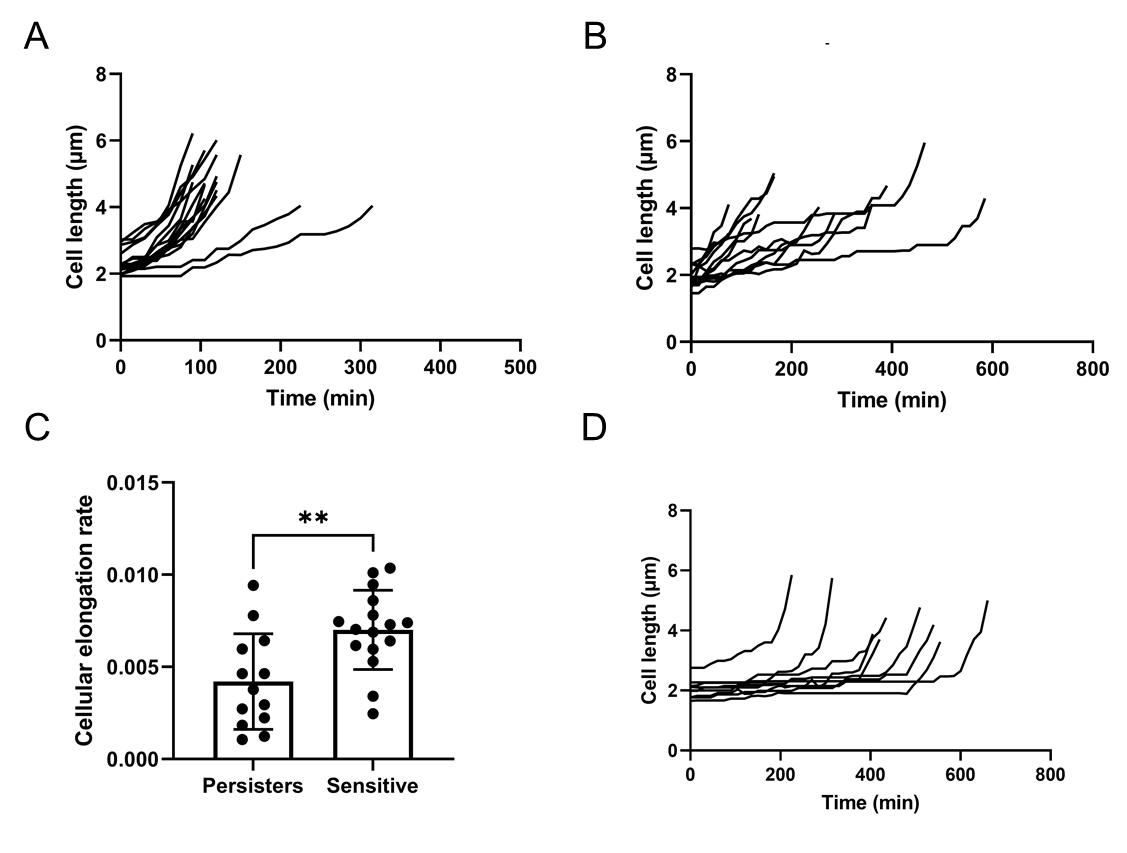

Figure S2. Cellular elongation rate during the awakening lag. Related to Figure 2. (A) Cell length in time for antibiotic-sensitive cells (wild type) during the awakening lag. (B) Cell length in time for persister cells (wild type) during the awakening lag. (C) By plotting an exponential curve on the data presented in Fig. S2A and S2B, the cellular elongation rate (k) of the cells can be inferred. In the awakening lag, persister cells have a decreased cellular elongation rate compared to the sensitive cells. Data represent mean ± SD. Asterisks indicate a significant difference (** P < 0.01). (D) *UvrD-*persisters have an increased awakening lag due to a stalled initiation of growth.

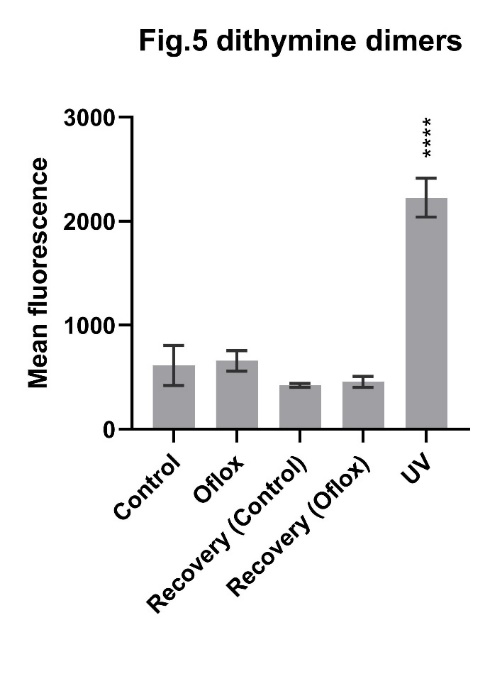

Figure S3. Ofloxacin treatment does not result in the formation of pyrimidine dimers, in contrast to UV-treatment. Related to Figure 3. Primary anti- pyrimidine antibodies were used, after which an Alexa Fluor secondary antibody, linked with a fluorescent protein, was used to quantify the formation of pyrimidine dimers. Fluorescence was measured using flow cytometry and the mean fluorescence of the population was calculated. Values result from n ≥ 3 biological repeats and are represented as mean ± SEM. Asterisks indicate significant difference (*** P < 0.001; ** P < 0.01).

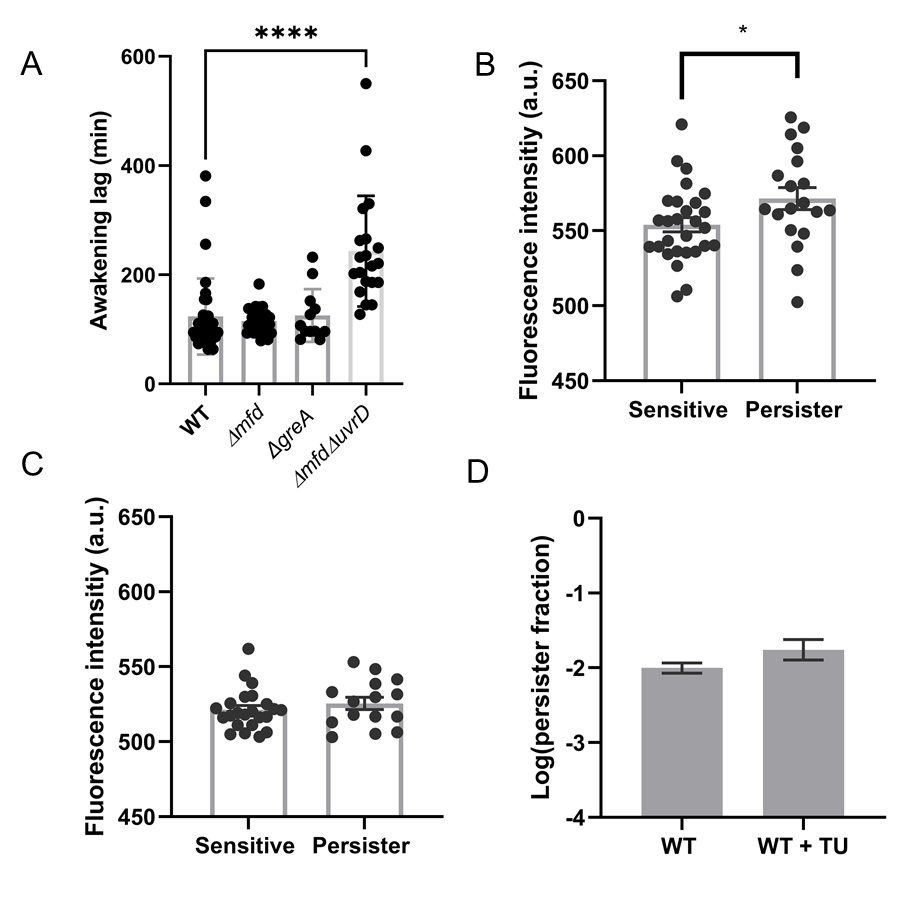

Figure S4. Alterations of the awakening lag in persister cells and sensitive cells. Related to Figure 4. (A) The awakening lag of *ΔmfdΔuvrD* sensitive cells following fluoroquinolone treatment differs significantly compared to the wild type. Values result from n ≥ 3 biological repeats and are represented as mean ± SD. Asterisks indicate a significant difference (**** P < 0.0001). (B) Initial fluorescence intensities following ofloxacin treatment in cells with the reporter plasmid Pmfd-gfp. A significantly increased expression is observed in persister cells compared to sensitive cells. Values result from n ≥ 3 biological repeats and are represented as mean ± SD. Asterisks indicate a significant difference. (C) Initial fluorescence intensities following ofloxacin treatment in cells with the reporter plasmid PuvrD-gfp. Values result from n ≥ 3 biological repeats and are represented as mean ± SD. (D) Addition of TU during ofloxacin treatment does not alter the number of surviving cells. The same culture was untreated or treated with TU. Values result from n ≥ 3 biological repeats and are represented as mean ± SEM.
